# A widespread Actinobacterial G Protein System regulates production of specialized metabolites in *Streptomyces coelicolor*

**DOI:** 10.1101/2025.09.26.678717

**Authors:** Luis M. Cantu Morin, Aditya Vunnum, Sydney Binns, Matthew F. Traxler

## Abstract

Actinobacterial G protein systems (AGPSs), also known as conservons, are regulatory systems that are broadly distributed within Actinomycetota. AGPSs are composed of a minimum of four proteins, including a sensor histidine kinase, a small Ras-like GTPase, a roadblock/MglB protein (likely a GTPase activating protein), and protein with a domain of unknown function that likely functions as a guanine-nucleotide exchange factor (GEF). While progress has been made in understanding AGPS function at the mechanistic level, the phylogenetic distribution of individual AGPSs, and the genes and processes they regulate, remain largely unmapped. Previously, the Cvn8 AGPS of *Streptomyces coelicolor* was found to influence expression of genes in multiple specialized metabolic pathways during interspecies interactions with other actinomycetes. However, the impact of the Cvn8 AGPS on specialized metabolism has not been assessed at the chemical level. Here, we investigated the phylogenetic distribution of the Cvn8 AGPS clade,and assessed the impact of the Cvn8 AGPS on natural product biosynthesis using untargeted metabolomics. In a set of 485 actinobacterial genomes, we found that members of the clade that includes the Cvn8 AGPS from *S. coelicolor* are widely distributed in the lineages known to produce specialized metabolites. We also found that in *S. coelicolor*, the pattern of specialized metabolite production varied in mutants lacking specific components of the Cvn8 AGPS. Specifically, normal production of the pigmented antibiotic actinorhodin during interspecies interactions required *cvnA8* and *cvnF8*, while a *ΔcvnD8* overproduced undecylprodigiosin. Together, these results connect a widespread AGPS to control of specialized metabolism in a model actinomycete.

**Importance:** Actinobacterial G protein systems (AGPSs) are found widely in bacteria in the phylum Actinomycetota, including genera like *Streptomyces* that produce many useful molecules and pathogens such as *Mycobacterium tuberculosis*. The genes and functions regulated by these regulatory systems are largely unknown. Here we investigated the role of the Cvn8 AGPS in controlling the production of specialized metabolites, like antibiotics, in the model actinomycete *Streptomyces coelicolor*. We found that specific components of the Cvn8 AGPS were required for normal production of the antibiotic actinorhodin during interactions between *S. coelicolor* and another actinomycete. We also show that Cvn8 belongs to a group of AGPSs that is found broadly in the genus *Streptomyces* and in more distantly related orders of Actinomycetota such as the Pseudonorcardiales and Micromonosporales. Together, these results raise the possibility that this group of AGPSs may influence specialized metabolism across a broad range of Actinomycetota lineages.

## Observation

Many organisms of the phylum Actinomycetota contain four component regulatory systems called Actinobacterial G protein systems (AGPSs), also known as conservons (1). AGPSs have been shown to control diverse functions, including fluoroquinolone resistance in pathogenic *Mycobacteria* (2–4) and aerial development and specialized metabolism in *Streptomyces* (5–7). Broadly speaking, signal transduction patterns within AGPSs are incompletely understood, as are the genes and processes that they regulate. Beyond this, the phylogenetic distribution of specific AGPS clades across Actinomycetota has yet to be explored.

Interspecies interactions with other actinomycetes can trigger production of specialized metabolites in the model organism *Streptomyces coelicolor* (8). We previously found that some components of the Cvn8 AGPS of *S. coelicolor*, including the accessory protein CvnF8, were required for visible induction of pigmented antibiotics when grown in proximity to the actinomycete *Amycolotopsis sp*. AA4 (9). Additionally, genes for production of multiple specialized metabolites including actinorhodins, prodiginines, coelimycins, and a cryptic lanthipeptide were differentially expressed in *cvn8* mutants (9). Notably, this differential expression occurred both during interspecies interactions, and when *cvn8* mutants were grown alone. However, the impact of the Cvn8 system (or any other AGPS) on specialized metabolism at the metabolomic level remains undefined. Here, we sought to use the Cvn8 AGPS of *S. coelicolor* as a model to characterize both the distribution of a specific AGPS clade and its function at the level of the metabolome.

*cvnA* homologs encode the histidine kinases that likely serve as sensory inputs for AGPSs. To map the evolutionary relationships of AGPSs, we built a tree using the *cvnA* sequences from 2,175 AGPSs (Fig. 1A) present in a collection of 485 Actinomycetota genomes (10). As noted previously, the Cvn8 and Cvn7 AGPSs of *S. coelicolor* belong to a larger clade of AGPSs that is defined by the absence of a nitrate sensing domain in their CvnA homologs, and the presence of a *cvnF* accessory gene (outlined in green and inset in Fig. 1A) (11). We suggest that AGPSs in this clade can be considered CvnF-dependent, as we previously showed that CvnF8 regulates the activity of the CvnA8 histidine kinase (11).

**Fig. 1.**
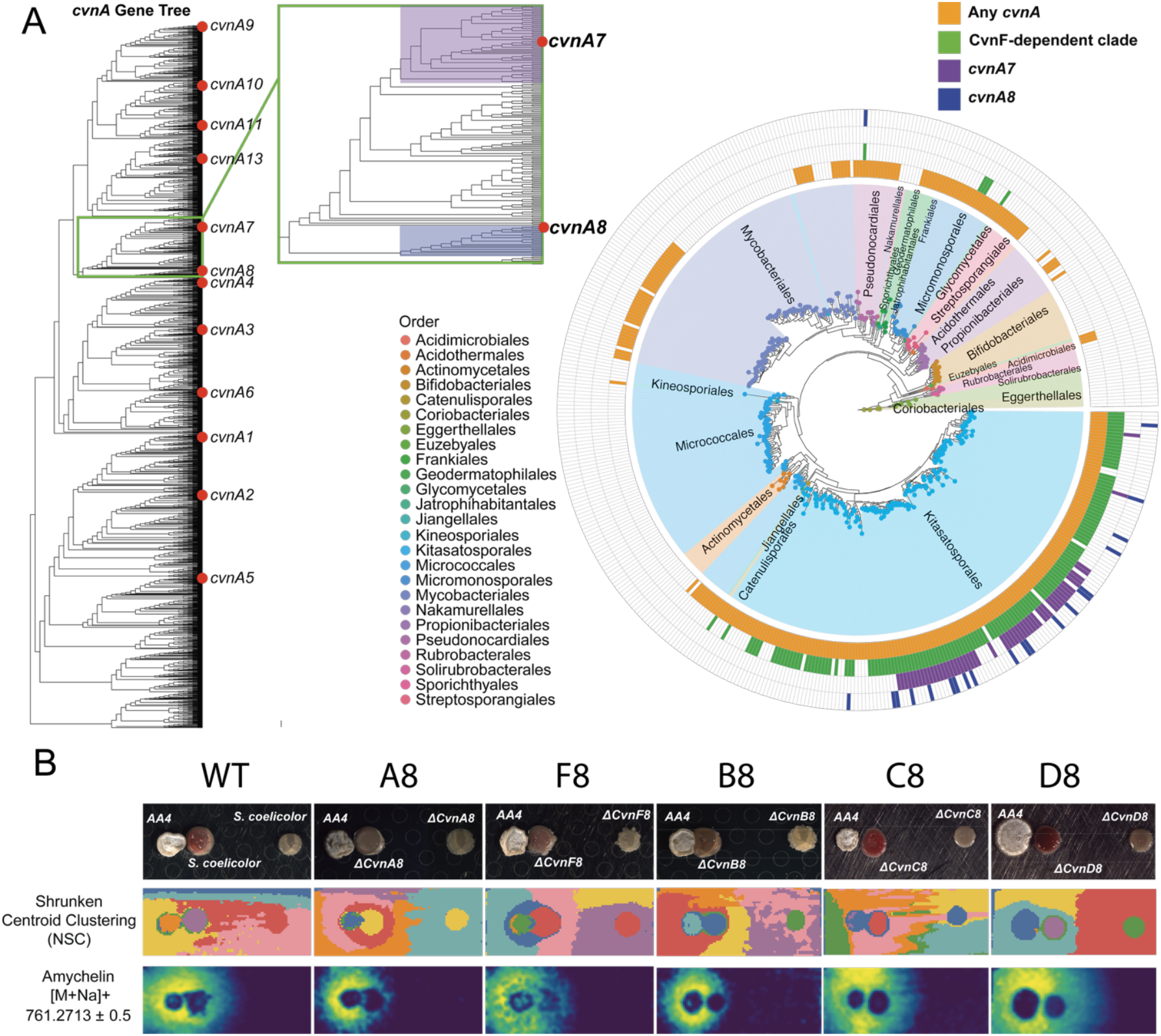
Phylogenetic distribution of the CvnA8 AGPS and its impact on spatial patterns of metabolite production. (A) Gene tree of *cvnA* orthologs identified by orthogrouping across 485 genomes from the ActDES Actinomycetota database. Red dots indicate proteins from *Streptomyces coelicolor*. The CvnF-dependent clade, which includes the *cvnA7* and *cvnA8* families, is highlighted (green box), and its distribution across species is shown in the circular species tree (right). Outer rings represent the presence of any *cvnA* (orange), the CvnF-dependent clade (green), *cvnA7* family (purple), and *cvnA8* family (blue). Species tree is colored by taxonomic order. (B) MALDI imaging mass spectrometry (MALDI-IMS) of *S. coelicolor cvnA8* deletion mutants during interactions with *Amycolatopsis* sp. AA4. Top row: colony morphology of wild-type and deletion strains. Middle row: Shrunken Centroid Clustering (NSC) shows distinct spatial metabolomic patterns. Bottom row: spatial distribution of amychelin (m/z 761.2713 ± 0.5), a siderophore produced by AA4 known to trigger chemical diversification. Loss of response in certain mutants suggests a role for Cvn proteins in mediating interaction-specific metabolic changes.

To understand the phylogenetic distribution of the Cvn8 and Cvn7 families, and the whole CvnF-dependent clade of AGPSs, we mapped their presence/absence across the Actinomycetota. Of the 485 genomes included, 291 contained at least one AGPS (orange track, Fig. 1A). Greater than ⅔ of the genomes with AGPSs contained a member of the CvnF-dependent clade (187 genomes, green track in Fig. 1A), while the individual Cvn7 and Cvn8 families were found in 56 and 22, genomes respectively. Representatives of the Cvn7 (purple track, Fig. 1A) and Cvn8 (dark blue track, Fig. 1A) families were found commonly in genomes of the order Kitasatosporales, which includes the genus *Streptomyces*, and in at least one member of the Pseudonocardiales. Additionally, other cryptic CvnF-dependent AGPSs were present in multiple organisms of the order Micromonosporales. Taken together, these analyses indicate that CvnF-dependent AGPSs (including the Cvn8 family) are found across multiple orders of Actinomycetota, with prevalence in lineages known for producing specialized metabolites.

We next investigated whether or not the Cvn8 AGPS influenced production of metabolites in *S. coelicolor* when grown alone, or in interactions with another actinomycete. For these experiments, we deposited patches of spores of wildtype *S. coelicolor*, and mutants lacking individual components of the Cvn8 system (*ΔcvnA8, ΔcvnB8, ΔcvnC8, ΔcvnD8*, and *ΔcvnF8* strains) in proximity to patches of *Amycolatopsis sp*. AA4 (3 mm away). We also placed another patch (the ‘alone’ patches) of the *S. coelicolor* strains farther away (2 cm) from the *Amycolatopsis sp*. AA4 patches, where they were visibly unaffected. The spores were allowed to germinate and grow for 72 hours.

To get a broad picture of the spatial patterns of metabolites produced by the *S. coelicolor* strains during interspecies interactions, we employed imaging mass spectrometry (12) and shrunken centroid clustering to identify chemically distinct spatial regions within the samples (13). Segmentation of the spatial data into eight clusters showed clear patterns in which the chemical signature of the wildtype patch in proximity to the *Amycolaptosis* sp. AA4 patch was distinct from the patch grown alone (leftmost panel, Fig. 1B). This pattern was also true of the *ΔcvnB8, ΔcvnC8*, and *ΔcvnD8* strains. In contrast, this pattern was disrupted in the patches of the *ΔcvnA8* and *ΔcvnF8* strains, which were not classified as chemically distinct even when grown in proximity to the *Amycolaptosis* sp. AA4 patches. We also observed that amychelin, a siderophore produced by *Amycolaptosis* sp. AA4 that is known to alter the specialized metabolite profile of *S. coelicolor*, was robustly produced under these conditions.

To obtain more detailed profiles of metabolites differentially produced depending on the interaction or mutant genotypes, we excised and extracted individual *S. coelicolor* patches in triplicate and performed an untargeted LC/MS analysis. A principal component analysis of the profiles obtained from all strains when grown alone (Fig. 2A), showed that all the mutant strain profiles grouped separately from the WT on the PC1 axis, which accounted for 40.9% of the observed variance. In contrast, during interactions, the pattern of metabolites produced by the *ΔcvnB8* and *ΔcvnC8* strains closely matched the wildtype, while those of the *ΔcvnA8, ΔcvnF8, and ΔcvnD8* diverged significantly across the first two principal components.

**Fig. 2.**
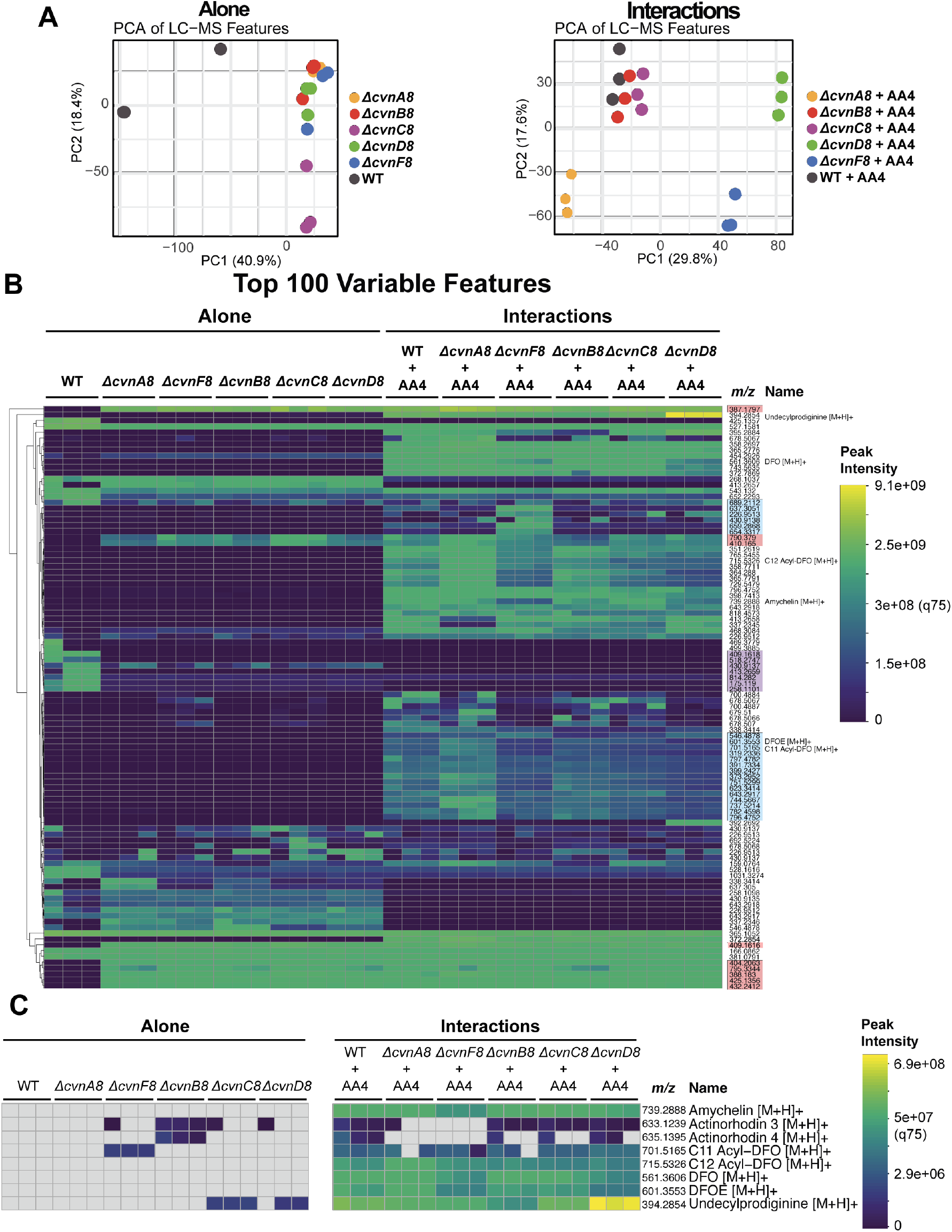
Metabolomic profiling of *cvnA8* clade mutants reveals altered chemistry during interspecies interactions. (A) Principal Component Analysis (PCA) of LC-MS features from methanol extracts of agar-grown colony patches. Mutants in the *cvnA8* clade form distinct clusters from wild-type *S. coelicolor*, both when grown alone and during interactions with *Amycolatopsis* AA4. (B) Heatmap of the top 100 most variable metabolite features across all samples, showing widespread differences in metabolic profiles. Several known specialized metabolites are annotated. (C) Focused heatmap of selected known compounds, including amychelin and actinorhodin derivatives. Disrupted or diminished production in mutants suggests a role for *cvn* genes in modulating specialized metabolism during interaction.

We next built a hierarchically-clustered heatmap composed of the top 100 variable features across all samples (Fig. 2B). Broadscale differences in metabolite patterns were obvious between patches that were grown alone and patches during interactions, with interacting patches producing a wider array of chemical features. In patches grown alone, we noted sets of features that were produced by the wildtype but not the mutants (*m/z*’s highlighted in purple), and features that were produced by the mutants but not the wildtype (highlighted in red). These results, combined with the PCA in Fig. 2A, indicate that *cvn8* mutants have a different metabolomic profile even when grown alone. During interactions, all of the features highlighted in red (which were constitutively made by the *cvn8* mutants) were made by all the strains, indicating that the wildtype activated the production of these metabolites in an interaction-dependent manner. Additionally, several groups of features were made at variable levels during interactions depending on the specific *cvn8* mutation (highlighted in light blue).

We next asked if specific known compounds were made differentially by the *cvn8* mutants (MS2 annotation provided in supplementary Figures S1-S8). We found that during interactions, the wildtype produced detectable amounts of two analogs of the antibiotic actinorhodin (Fig. 2C), but these were largely undetectable in the *ΔcvnA8* and *ΔcvnF8* strains. Additionally, the *ΔcvnD8* strain produced higher levels of the red antibiotic undecylprodiginine. These trends aligned with our previous findings that during interactions, *ΔcvnA8* and *ΔcvnF8* strains expressed genes of the actinorhodin pathway at significantly lower levels, while the *ΔcvnD8* strain showed enhanced expression of the pathway for prodiginine biosynthesis (9).

Taken together, the results presented here demonstrate that the Cvn8 AGPS plays a key role for normal production of specialized metabolites in *S. coelicolor*, both alone and during interspecies interactions. These metabolomic results, combined with the phylogenetic distribution of the Cvn8 AGPS and related systems, opens the possibility that these AGPSs may influence specialized metabolism across many lineages of Actinomycetota.

## Acknowledgements

L. C. acknowledges support from the National Science Foundation Graduate Research Fellowship Program. Funding for this work was provided by NIH R35GM128849 awarded to M. F. T.

